# A universal reporter cell line for bioactivity evaluation of engineered cytokine products

**DOI:** 10.1101/739987

**Authors:** Jacqueline Mock, Christian Pellegrino, Dario Neri

**Affiliations:** Department of Chemistry and Applied Biosciences, Swiss Federal Institute of Technology (ETH Zürich), Vladimir-Prelog-Weg 4, CH-8093 Zürich (Switzerland)

## Abstract

Engineered cytokine products represent a growing class of therapeutic proteins which need to be tested for biological activity at various stages of pharmaceutical development. In most cases, dedicated biological assays are established for different products, in a process that can be time-consuming and cumbersome. Here we describe the development and implementation of a universal cell-based reporter system for various classes of immunomodulatory proteins. The novel system capitalizes on the fact that the signaling of various types of pro-inflammatory agents (e.g., cytokines, chemokines, Toll-like receptor agonists) may involve transcriptional activation by NF-κB. Using viral transduction, we generated stably-transformed cell lines of B or T lymphocyte origin and compared the new reporter cell lines with conventional bioassays. The experimental findings with various interleukins and with members of the TNF superfamily revealed that the newly-developed “universal” bioassay method yielded bioactivity data which were comparable to the ones obtained with dedicated conventional methods. The engineered cell lines with reporters for NF-κB were tested with several antibody-cytokine fusions and may be generally useful for the characterization of novel immunomodulatory products. The newly developed methodology also revealed a mechanism for cytokine potentiation, based on the antibody-mediated clustering of TNF superfamily members on tumor-associated extracellular matrix components.

## INTRODUCTION

The clinical success of immune check-point inhibitors for the treatment of various forms of cancer ^1-3^ has sparked research activities for the discovery and development of novel immunostimulatory products. Various types of engineered cytokine products (e.g., antibody-cytokine fusion proteins ^4-6^ and polymer-cytokine conjugates ^7,8^, chemokines ^9^ and Toll-like receptor agonists ^10^) have been considered for cancer therapy applications.

The development of novel immunostimulatory products requires the implementation of reliable quantitative methods for the determination of biological activity. Such methods are important both at the research stage and during industrial production, since a demonstration of consistent biological activity throughout the development process is a prerequisite for quality assurance and for enabling comparisons of experimental results.

At present, dedicated assays are established for individual products but most methodologies are not readily applicable to different types of biopharmaceuticals^11^. For example, interleukin-2 (IL2) activity is typically measured by the ability to induce proliferation of the CTLL-2 cell line of T cell origin^12^, while Interleukin-12 (IL12) activity tests measure the production of interferon-*γ* by NK-92 cells of NK cell origin ^13,14^. The activity of tumor necrosis factor (TNF) and of TNF-based biopharmaceuticals is often measured by the killing of transformed fibroblast cell lines ^15^, while other members of the TNF superfamily are studied in terms of their ability to stimulate the production of pro-inflammatory cytokines by splenocytes ^16,17^ or the proliferation of splenic subsets ^18,19^.

The development of dedicated methods for individual products is often cumbersome and may require the generation of stably transfected cell lines ^20^. Moreover, the optimization of technical parameters, such as cell growth conditions and dose-response relationships, may be time-consuming. Our laboratory has worked on the development and characterization of more than 100 different antibody-cytokine and antibody-chemokine fusions proteins ^4,21^ and has learned to value the importance of robust, reliable and broadly applicable methodologies for the study of engineered cytokine products.

Here we report on the development of two general reporter cell lines, which are derived from T and B lymphocytes and which are broadly applicable for the quantification of biological activity of immunostimulatory agents. We used viral transduction methodologies with a reporter for NF-κB activity because of the central role played by NF-κB signaling in many different inflammatory processes ^22^.

The term NF-κB refers to a variety of homo- and heterodimers that are formed between members of the NF-κB family of proteins. The members of the NF-κB family of proteins all share a related REL homology domain (RHD), which confers both DNA binding and dimerization. Activated NF-κB dimers localize to the nucleus where they bind to the NF-κB response element that has the loose consensus sequence 5′-GGRNN(WYYCC)-3′ (where R: purine, N: any base, W: adenine or thymine and Y: pyrimidine) ^23-25^ and thus activates the transcription of a variety of target genes (via interaction with basal transcription factors and cofactors). In the absence of signaling, the NF-κB proteins are present in the cell as pre-formed complexes that can be rapidly activated upon signaling ^24,25^. The signaling complexes can either be activated by degradation of an inhibitory protein or by the removal of an inhibitory protein domain by upstream signaling components ^26^.

Even though class I cytokines typically signal via JAK/STAT activation ^27^, there is evidence that most of them directly or indirectly also trigger the activation of NF-κB ^28-31^. By contrast, members of the TNF superfamily of proteins signal via the recruitment of TNF Receptor Associated Factors (TRAF) that activate NF-κB both via the classical and the non-canonical pathway ^32-34^.

The newly developed cell lines (termed CTLL-2_NF-κB and A20_NF-κB) were used to implement a general bioassay, which was compared to established procedures for the characterization of various types of engineered cytokine products. Moreover, the new methodologies allowed the discovery of a novel strategy for cytokine potentiation, based on the antibody-mediated clustering of multimeric immunostimulatory payloads (e.g., members of the TNF superfamily) on tumor-associated extracellular matrix components.

## MATERIALS AND METHODS

### Cell lines

The murine cytotoxic T cell line CTLL-2 (ATCC® TIB-214), the murine B lymphocyte cell line A20 (ATCC® TIB-208) and the murine fibroblast cell line LM (ATCC® CCL-1.2) and were obtained from ATCC, expanded and stored as cryopreserved aliquots in liquid nitrogen. The CTLL-2 cells were grown in RPMI-1640 (Gibco, #21875034) supplemented with 10% FBS (Gibco, #10270106), 1 X antibiotic-antimycoticum (Gibco, #15240062), 2 mM ultraglutamine (Lonza, #BE17-605E/U1), 25 mM HEPES (Gibco, #15630080), 50 μM β-mercaptoethanol (Sigma Aldrich) and 60 U/mL human IL-2 (Proleukin, Roche Diagnostics). The A20 cells were grown in RPMI-1640 (Gibco, #21875034) supplemented with 10% FBS (Gibco, #10270106), 1 X antibiotic-antimycoticum (Gibco, # 15240062) and 50 μM β-mercaptoethanol (Sigma Aldrich). The L-M fibroblasts were grown in DMEM (Gibco, high glucose, pyruvate, #41966-029) supplemented with 10% FBS (Gibco, #10270106) and 1 X antibiotic-antimycoticum (Gibco, #15240062). The cells were passaged at the recommended ratios and never kept in culture for more than one month. The NK-92 cells were obtained from DSMZ (ACC 488) and grown in in MEM Alpha medium (Gibco, #22571-020) supplemented with 5 ng/mL recombinant human interleukin 2 (Gibco, #PHC0027), 2 mM L-glutamine (Lonza, #17-605E), 12.5% Fetal Bovine Serum (Gibco, #10099-141) and 12.5% Horse serum (Sigma Aldrich, #H1270).

### Cloning of the reporter construct

The plasmid pNL3.2.NF-κB-RE[NlucP/NF-κB-RE/Hygro] (#N1111) encoding the NanoLuc luciferase under the control of the NF-κB response element was obtained from Promega AG. The IL-6 secretion signal was inserted into the plasmid by polymerase chain reaction (PCR) and subsequent blunt-end ligation. The cassette was then transferred into a lentiviral transfer vector by Gibson Isothermal assembly. The lentiviral vector was based on the EF1-T2A vector which was kindly provided by Dr. Renier Myburgh and Prof. Dr. Markus Manz. A detailed representation of the vector pJM046 and the DNA sequence can be found in [**Supplementary Figures S1 and S2**].

### Protein production

Various antibody-cytokine fusions were produced and tested in this work. The cloning and construction of L19-IL2 ^35^, F8-TNF ^36^ and L19-IL12 ^37^ is described elsewhere. Soluble single-chain trimers of 4-1BBL, glucocorticoid-induced tumor necrosis factor receptor ligand (GITRL) and CD154 were designed by linking the extracellular domain with suitable glycine-serine linkers. Genetic sequences encoding the TNF-homology domain of murine 4-1BBL (amino acids 139 – 309), of the extracellular domain of murine GITRL (amino acids 46 – 170) and of the soluble part of murine CD154 (amino acids 112 – 260) as single-chain trimers were ordered from Eurofins Genomics. These sequences were then introduced into vectors encoding the F8 in a diabody format by Gibson Isothermal Assembly. To clone the single-chain variable Fragment (scFv) linked to the TNFSF monomer, the genetic sequence encoding the diabody was replaced by the sequence encoding the scFv and two domains of 4-1BBL and GITRL respectively were removed by PCR followed by blunt-end ligation. The protein sequences are provided in [**Supplementary Figure S3**].

Proteins were produced by transient transfection of CHO-S cells and purified by protein A affinity chromatography as described previously ^35-37^. Quality control of the purified products included SDS-PAGE, size exclusion chromatography and, where applicable, mass spectrometry [**Supplementary Figures S4 and S5**].

### Virus production and stable transduction

For the virus production, 5 million HEK293T cells were seeded at a density of 300,000 cells/mL on the day prior to transfection. They were then transiently co-transfected with the reporter plasmid pJM046 as well as the packaging plasmid psPAX2 (Addgene, #12260; http://n2t.net/addgene:12260; RRID:Addgene_12260) and the envelope plasmid pCAG-VSVG (Addgene, #35616; http://n2t.net/addgene:35616; RRID:Addgene_35616) using the jetPRIME reagent (Polyplus transfection). The packaging and the envelope plasmid were kindly provided by the group of Prof. Dr. Patrick Salmon. The medium was replaced on the day after the transfection and the virus was harvested on the following day. The virus was aliquoted and snap-frozen in an ethanol dry ice mixture. An aliquot was thawed and used for the transduction of 500,000 target cells. For the transduction 500,000 cells in 1 mL of medium were seeded in a 24 well plate and 1 mL of virus was added. Polybrene (Santa Cruz Biotechnology, #134220) was added to a final concentration of 8 μg/mL. The cells were then centrifuged for 90 min at 1000 × g, 32°C. Afterwards, the cells were incubated at 37°C, 5% CO_2_. The medium was exchanged daily for the following three days. The cells were then expanded, activated and mCherry-positive cells were sorted by FACS (BD FACS AriaIII).

### CTLL-2 proliferation assay

In order to starve the CTLL-2 cells from IL-2, the cells were washed twice with prewarmed HBSS (Gibco, #14175095) and then grown in the absence of IL-2 for 24 h in RPMI-1640 (Gibco, # 21875034) medium supplemented with 10% FBS (Gibco, #10270106), 1 X antibiotic-antimycoticum (Gibco, # 15240062), 2 mM ultraglutamine (Lonza, # BE17-605E/U1), 25 mM HEPES (Gibco, # 15630080) and 50 μM β-mercaptoethanol (Sigma Aldrich). The starved CTLL-2 cells were seeded in a 96-well plate (20’000 cells/well) and medium supplemented with varying concentrations of IL-2 was added. The total volume per well was 200 μL which corresponds to a concentration of 100,000 cells/mL. All dilutions were done in triplicates. After 48 h incubation at 37°C, 5% CO_2_ CellTiter 96 Aqueous One Solution (Promega, #G3582) was added to measure cell proliferation. Absorbance at 490 nm and 620 nm was measured after 1.5 h. The relative proliferation was calculated using the formula: relative proliferation = (OD_490-620_^treated^-OD_490-620_^medium^)/ (OD_490-620_^untreated^-OD_490-620_^medium^) × 100%. The data was fitted using the [Agonist] vs. response (three parameters) fit of the GraphPad Prism 7.0 a software to estimate the EC_50_.

### IL-12 assay

The method was adapted from previous publications^14,38^. Briefly, NK92 cells were seeded at a density of 100,000 cells per well in a 96-well plate and 100 μL of medium containing varying concentrations of L19-IL12 was added. The total volume per well was 200 μL which corresponds to a cell density of 500,000 cells/mL. After 24 h of incubation at 37°C, 5% CO_2_, the concentration of IFN-γ in the cell culture supernatant was determined by ELISA (Invitrogen, #EHIFNG2). An IFN-γ standard was included in the ELISA and linear curve fitting using the GraphPad Prism 7.0 a software was used to derive the IFN-γ concentration in the samples from the absorbance at 450 nm (A450) **[Supplementary Figure S7]**. The data was fitted using the [Agonist] vs. response (three parameters) fit of the GraphPad Prism 7.0 a software to estimate the EC_50_.

### TNF assay

L-M fibroblasts were seeded at a density of 20,000 cells/well in a 96-well plate and incubated for 24 h at 37°C, 5% CO_2_. The medium was replaced by 100 μL fresh medium containing 2 μg/mL actinomycin D (BioChemica, #A1489,0005) and varying concentrations of F8-TNF. The cells were seeded at a concentration of 100,000 cells/mL. One day after seeding, the cell density should correspond to approximately 200,000 cells/mL, in a total of 200 µL, but this concentration was not measured immediately prior to the assay. The cells were incubated at 37°C, 5% CO_2_ for another 24 h before 20 μL of CellTiter 96 Aqueous One Solution (Promega, #G3582) was added and absorbance at 490 nm and 620 nm was measured. The % cell survival was calculated using the formula: relative proliferation = (OD_490-620_^treated^-OD_490-620_ ^medium^)/ (OD_490-620_^untreated^-OD_490-620_ ^medium^) × 100%. The data was fitted using the [Agonist] vs. response (three parameters) fit of the GraphPad Prism 7.0 a software to estimate the EC_50_.

### NF-κB response assay

CTLL-2 reporter cells were starved for 6 – 9 h as described above prior to use in order to reduce the background signal. If necessary, 100 μL 100 nM 11-A-12 fibronectin in phosphate buffered saline (PBS) was added to each well to be coated with EDA and the plate was incubated at 37°C for 90 min. Cells were seeded in 96-well plates (50,000 cells/well) and growth medium containing varying concentrations of the cytokine to be tested was added. The cells were incubated at 37°C, 5% CO_2_ for several hours. To assess luciferase production, 20 μL of the supernatant was transferred to an opaque 96-well plate (PerkinElmer, Optiplate-96, white, #6005290) and 80 μL 1 μg/mL Coelenterazine (Carl Roth AG, #4094.3) in phosphate buffered saline (PBS) was added. Luminescence at 595 nm was measured immediately. mCherry expression was measured by flow cytometry (CytoFLEX, Beckman Coulter) and the data was analyzed using FlowJo (v.10, Tree Star). The cells were resuspended in growth medium and transferred to a 96-well U bottom plate (Greiner BioOne, Cellstar, #650180) and harvested by centrifugation. The medium was discarded and the cells were washed with FACS buffer (0.5% BSA, 2 mM EDTA, PBS) and resuspended in FACS buffer. The relative luminescence and the relative fluorescence were calculated by dividing the obtained results by the results obtained when no inducer was added.

## RESULTS

**Figure 1** describes the strategy followed for the generation of a universal reporter system for cytokine activity. The signaling of many different pro-inflammatory mediators (e.g., chemokines, Toll-like receptor agonists, members of the TNF superfamily and various other cytokines) involves the activation of NF-κB, in addition to other signaling pathways [**Figure 1a**]^23,30,39,40^. We constructed a vector for virus-mediated stable cell transduction, which incorporated an NF-κB response element upstream of a secreted luciferase (NanoLuc) and mCherry reporter genes [**Supplementary Figures S1 and S2**]. The two reporter proteins are separated by a T2A peptide. The luciferase is secreted due to the presence of an IL-6 secretion signal while the mCherry is retained in the cytoplasm. A second-generation lentivirus was used for transduction experiments. The lentivirus was produced by the simultaneous transfection of HEK293T cells with the envelope plasmid pCAG-VSVG, the packaging plasmid psPAX2 and pJM046 (harboring the reporter construct) [**Supplementary Figures S1 & S2**]. The transduction strategy was used to stably integrate the NF-κB reporter into CTLL-2 and A20 cell lines, of T-cell and B-cell origin (respectively) [**Figure1**].

**Figure 1:**
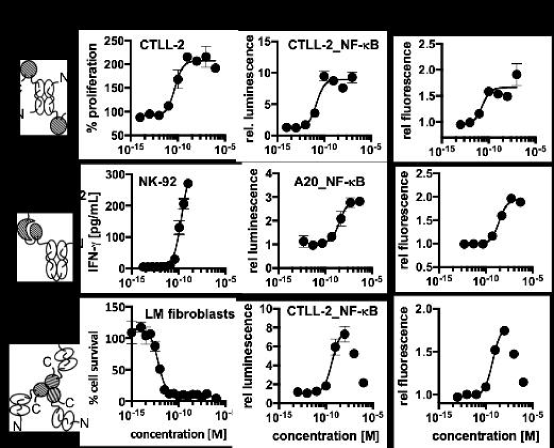
General strategy for the design of the reporter cell line: **[a]** signal through most of the immunologically relevant receptors results in the activation of NF-κB ^23,30,39,40^ **[b]** the reporter construct encoding a secreted luciferase (NanoLuc) and mCherry separated by a T2A sequence downstream of an NF-κB response element (NF-κB-RE) was used for the viral transduction of cell lines of T and B cell origin (TCR: T cell receptor, TNFR: TNF receptor, IL4 R: IL-4 receptor, IL2 R: IL-2 receptor, BCR: B cell receptor, IL12 R: IL-12 receptor, TLR: Toll like receptor)

We investigated the possibility of using the newly developed transduction method as a universal reporter for cytokine activity by performing a comparison with established dedicated test systems. For this experiment, antibody-cytokine fusions that have previously been developed in our lab were used ^35-37^. We used fusions of the L19 antibody targeting the extradomain B of fibronectin ^41^ linked to human IL2 and murine IL12, as well as fusions of the F8 antibody targeting the extradomain A of fibronectin ^42^ linked to murine tumor necrosis factor (TNF) [**Figure 2**]. Interleukin-2 (IL2) activity was measured both in terms of proliferation of non-transduced CTLL-2 cells and in terms of NF-κB reporter activity in transduced cells. An EC_50_ value in the 10 pM range was observed for both methodologies [**Figure 2a**] [**Supplementary Table T1**]. The activity of interleukin-12 (IL12) is often measured in terms of interferon-γ production by NK-92 cells of Natural Killer cell origin^14^. In this case, the EC_50_ values obtained using the NK-92 cell-based system were roughly ten times lower than those obtained by monitoring NF-κB reporter activity [**Figure 2b** and **Supplementary Table T1**]. The performance of the new methodology for the measurement of TNF activity was compared to the results of a conventional cell killing assay, using a transformed fibroblast cell line which is particularly sensitive to the action of TNF. EC_50_ values in the pM range were observed for the killing assay, whereas the EC_50_ values obtained with the universal assay were in the 100 pM range [**Figure 2C** and **Supplementary Table T1**].

**Figure 2:**
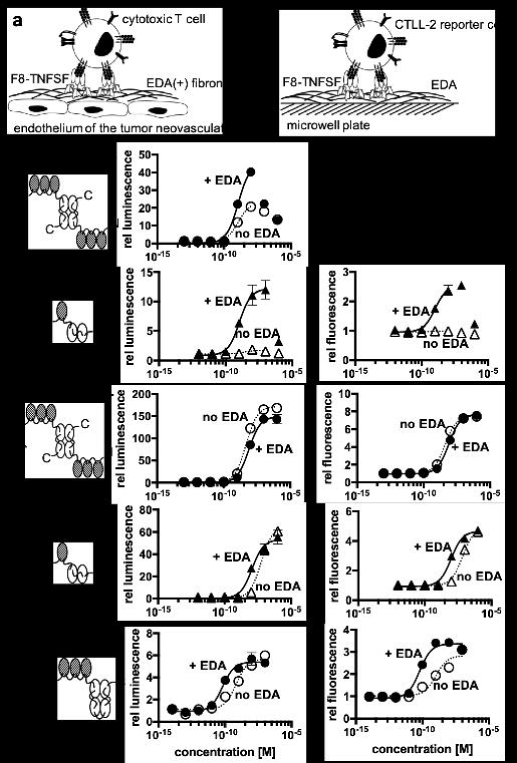
Comparison of the expression of luciferase and mCherry by the newly generated reporter cell lines with the response obtained with the same proteins in the conventional assay. The immunocytokines that were used for this assay are schematically depicted (the antibody moiety is depicted as empty symbols and the cytokine moieties as hatched areas) **[a]** the proliferation of CTLL-2 and the response of the CTLL-2 reporter cell line triggered by L19-IL2 **[b]** the production of interferon-γ by NK-92 cells and the response of the A20 reporter cell line triggered by L19-IL12 **[c]** the cytotoxicity of F8-TNF for L-M fibroblasts compared to the response of the CTLL-2 reporter cell line to F8-TNF. In the cases where a strong hook effect was observed, only the sigmoidal part was used for curve fitting as indicated by the solid line.

We then used the of NF-κB reporter assay for the characterization of three novel fusion proteins, featuring the F8 antibody (specific to the alternatively-spliced EDA domain of fibronectin, a tumor-associated antigen) ^42^ fused to three different members of the TNF superfamily [**Figure 3**]. 4-1BB and GITR (glucocorticoid-induced TNFR-related gene) are expressed on activated CD8^+^ T-cells. Activation of each of these two receptors has been shown to prolong longevity of the activated T cells and transition towards a memory phenotype^43,44^. By contrast, CD40 is expressed on antigen-presenting cells^45^. The interaction between CD40 and its ligand, CD154, is important for the licensing of antigen-presenting cells^46^. A variety of agonists to these three members of the TNF superfamily are being developed and tested both at a preclinical stage and in clinical trials as agents for the immunotherapy of cancer^47^. Our lab has had a long-standing interest in the characterization of the tumor-homing properties ^48^ and anti-cancer activity of antibody fusion proteins with members of the TNF superfamily ^36,49^. F8-41BBL, F8-GITRL and F8-CD154 featured the TNF superfamily member as single-chain polypeptide in order to stabilize the homotrimeric structure ^20^ while the antibodies were used as recombinant diabody moieties ^50^. A concentration-dependent NF-κB reporter activity could be measured for all three antibody-cytokine fusions, monitoring both luciferase activity and mCherry expression [**Figure 3**]. The EC_50_ values obtained for 4-1BBL and GITRL constructs were in the nanomolar range whereas the values obtained for the CD154 construct were in the 100 pM range [**Supplementary Table T2**].

**Figure 3:**
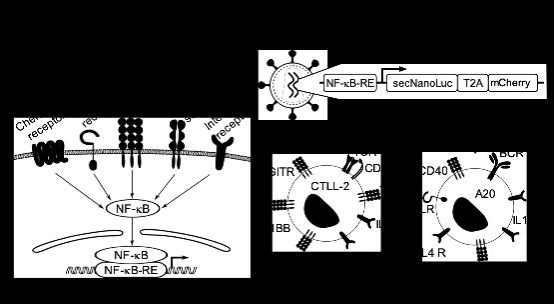
Activity tests of novel antibody-cytokine conjugates using the newly developed reporter cell lines **[a]** activity tests of the single chain trimeric 4-1BBL fused to the F8 antibody in the diabody format using the CTLL-2 reporter cell line **[b]** activity of single chain trimeric GITRL fused to the F8 antibody in the diabody format using the CTLL-2 reporter cell line **[c]** activity of single-chain trimeric CD154 fused to the F8 antibody in the single-chain diabody format using the A20 reporter cell line. In the cases where a strong hook effect was observed, only the sigmoidal part was used for curve fitting as indicated by the solid line.

Antibody-cytokine fusions are increasingly being used for the therapy of cancer ^5,6,21^ and of chronic inflammatory conditions ^51,52^ with the aim to concentrate cytokine activity at the site of disease and help spare normal organs. We used derivatives of the F8 antibody to assess whether a localized concentration of cytokine activity in close proximity to target cells of interest may lead to a potentiation of biological activity. The alternatively-spliced extra-domain A (EDA) of fibronectin (i.e., the target antigen of the F8 antibody) is typically found as an abundant component of the modified extracellular matrix associated with newly-formed tumor blood vessels ^53^, but is otherwise undetectable in most normal adult tissues, exception made for the female reproductive system ^54^. We mimicked the localized deposition of EDA(+)-fibronectin by coating plastic wells with 11-A-12 recombinant fibronectin fragments, containing the EDA domain ^42^ [**Figure 4a**]. The system was used to compare the biological activity of F8-41BBL, F8-GITRL and F8-CD154 fusion proteins, featuring the F8 antibody either in diabody format linked to a single-chain trimer of the cytokine or as single-chain Fv fragment linked to a single unit of the cytokine [**Figure 4b-d**]. In the case of the F8 antibody in a diabody format linked to a single-chain trimeric 4-1BBL, only a small increase in stimulatory activity was observed when the agonist was clustered on EDA. However, in the case of the single-chain Fv fragment linked to a single 4-1BBL unit, no activity was observed in the absence of clustering. Activity could in this case be restored by clustering the agonist on EDA coated wells [**Figure 4b**]. By contrast, both F8-GITRL constructs were constitutively active [**Figure 4c**]. In the case of the F8 antibody in a diabody format linked to the single-chain trimer of GITRL, no activity enhancement was observed by clustering on EDA. The EC_50_ of the construct consisting of the F8 antibody in an scFv format linked to a single unit of GITRL, a roughly 5-fold reduction in the EC_50_ was observed when the experiment was performed in the presence of EDA. Some modest increase in activity in the presence of EDA was also observed in the case of the single-chain trimeric CD154 linked to F8 in a single-chain diabody format [**Figure 4d**].

**Figure 4:**
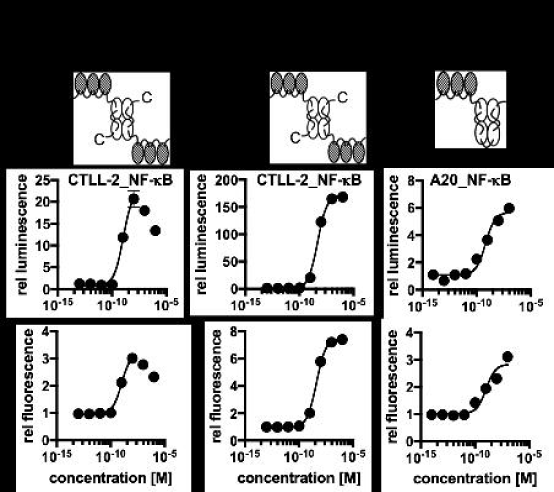
Clustering of cytokines of the TNF superfamily fused to fragments of the F8 antibody on EDA **[a]** *In vivo* the F8 antibody binds to the extradomain A (EDA) of fibronectin that is present in the tumor-associated neovasculature. This is thought to lead to clustering of the cytokine fused to the F8 antibody and therefore to enhance activation of target cells such as cytotoxic T cells. To mimic this situation *in vitro*, microwell plates were coated with the extradomain A of fibronectin. The activity of the immunocytokine when clustered on EDA and without clustering was compared for different formats. **[b]** activity of 4-1BBL in the single-chain trimeric format fused to the F8 antibody in the diabody format and in the monomeric format fused to the single-chain Fragment variable of F8 (scFv) **[c]** activity of GITRL in the single-chain trimeric format fused to the F8 antibody in the diabody format and in the monomeric format fused to the single-chain Fragment variable of F8 **[d|** activity of CD154 in the single-chain trimeric format fused to the F8 antibody in the single chain diabody format (circles: F8 in diabody format, triangles: F8 in scFv format, filled symbols: plate coated with EDA, empty symbols, dashed line: no EDA)

## DISCUSSION

In this work, we presented the development of two reporter cell lines that can be used to measure the biological activity of a variety of cytokines. The new methodology capitalizes on the fact that many cytokine-triggered signaling events converge at the level of NF-κB activation.

An excellent agreement could be found between bioactivity measurements of IL-2 performed with the conventional cell-based cytokine activity assay and the newly developed method [**Figure 2a**]. A discrepancy between the conventional cell-based cytokine assay and the new method could be seen for two fusion proteins with murine payloads (L19-IL12 and F8-TNF), for which the new methodology indicated a reduction in potency [**Figure 2b and 2c**]. It is possible that NF-κB signaling may not fully capture the molecular events triggered by murine IL-12 and murine TNF, but the new methodology could still enable a comparative evaluation of multiple pro-inflammatory payloads and cytokine variants (e.g., wildtype and mutated versions).

The observation that the activity of murine 4-1BBL can be potentiated by clustering of antibody fusions on specific tumor-associated extracellular matrix components is surprising and potentially useful for pharmaceutical applications. Murine 4-1BBL is not able to form stable homotrimers, but rather forms low-activity homodimeric structures^55,56^. It is possible that the high-density binding of F8 fusions on EDA-containing fibronectin promotes the formation of supermolecular assemblies, which gain signaling activity in a concentration-dependent manner [**Figure 4b**]. This approach mimics on the extracellular matrix what happens on murine cells where surface display of multiple copies of dimeric ligands turns an inactive homodimer into an active multimeric assembly^55,56^. Some potentiation upon antigen binding had previously been reported for certain murine TNF fusions^57^ and for other members of the TNF superfamily^58,59^. In principle, it would be attractive to engineer antibody-cytokine fusions which gain activity at the site of disease (e.g., upon antigen binding) while sparing normal tissues, as this approach could lead to biopharmaceuticals with improved therapeutic index. Our group has recently described a conceptually similar strategy, based on the assembly of heterodimeric split cytokine fusions (e.g., interleukin-12 superfamily members)^60^. Other strategies for the conditional potentiation of antibody-cytokine include the allosteric regulation of cytokine activity^61^ or the attenuation of cytokine potency (“attenukine”)^62^. Strategies aimed at potentiating the activity of antibody therapeutics based on conditional oligomerization include the development of hexameric IgG antibodies to augment complement-mediated cytotoxicity^63,64^.

Not all TNF superfamily members seem to need antigen-dependent clustering in order to gain activity [**Figure 2c and 4c**]. This is also reflected by the fact that some ligands of the TNF superfamily including TNF and CD154 are enzymatically shed *in vivo* and act as soluble ligands^65,66^. Although GITRL is so far not known to be present as soluble ligand^67^ our data indicates that the recombinant soluble ligand shows similar behavior [**Figure 4c**].

In this article we have mainly focused on recombinant and engineered cytokine products. While many interleukins mainly signal through JAK/STAT activation ^27^, NF-κB-driven transcriptional events are also induced, in line with previous reports on this matter ^28-31^. NF-κB reporters should be broadly applicable also to other classes of pro-inflammatory products. Both Toll-like receptors and members of the interleukin-1 receptor superfamily activate NF-κB through the recruitment of MyD88 as part of their signal transduction ^39^. In addition, there is also evidence that NF-κB can be activated by chemokine signaling ^40^.

The CTLL-2 and A20 cell lines express on their surface a large variety of receptors of immunological importance and should be therefore applicable for many different bioactivity assays. If a researcher is interested in the use of a different cell line, the viral transduction system described in **Figure 1** should enable the rapid preparation of the corresponding reporter system. The intracellular expression of mCherry was found to be useful for sorting of positively transduced cells by flow cytometry, while the secreted luciferase rapid a facile readout of the reporter activity.

In summary, we have generated new universal reporter systems for the facile measurement of biological activity of various types of pro-inflammatory mediators and engineered cytokine products. We anticipate that the newly-developed cell lines and vector may find a broad applicability in biological, pharmaceutical and immunological research.

## ACKNOWLEDGEMENTS

Financial support by the ETH Zürich, the Swiss National Science Foundation (grant number 310030B_163479/1), the European Research Council (ERC) under the European Union’s Horizon 2020 research and innovation program (grant agreement 670603), and the Federal Commission for Technology and Innovation (KTI, grant number 12803.1 VOUCH-LS) is gratefully acknowledged. In addition, the *ETH Zurich Flow Cytometry Core Facility* is acknowledged for their help with the sorting of the transduced cell lines. We also thank scientists at Philogen for help with the development of some cytokine assays. Further, we thank Emanuele Puca for providing the samples of L19-IL2 and L19-IL12 that were used for this study. Moreover, we would like to thank Dr. Renier Myburgh and Prof. Dr. Markus Manz for kindly providing the lentiviral vectors.

## AUTHOR CONTRIBUTIONS

D.N. and J.M. designed and planned the study. J.M. carried out most of the experimental work and prepared the figures. C.P. was involved in the design of the reporter construct and the virus production. D.N. and J.M. wrote the manuscript.

## DATA AVAILABILITY STATEMENT

The datasets generated during and/or analyzed during the current study are available from the corresponding author on reasonable request.

