## SupplementaryData for "A universal reporter cell line for bioactivity evaluation of engineered cytokine products"

*) Corresponding Author

| **a** | 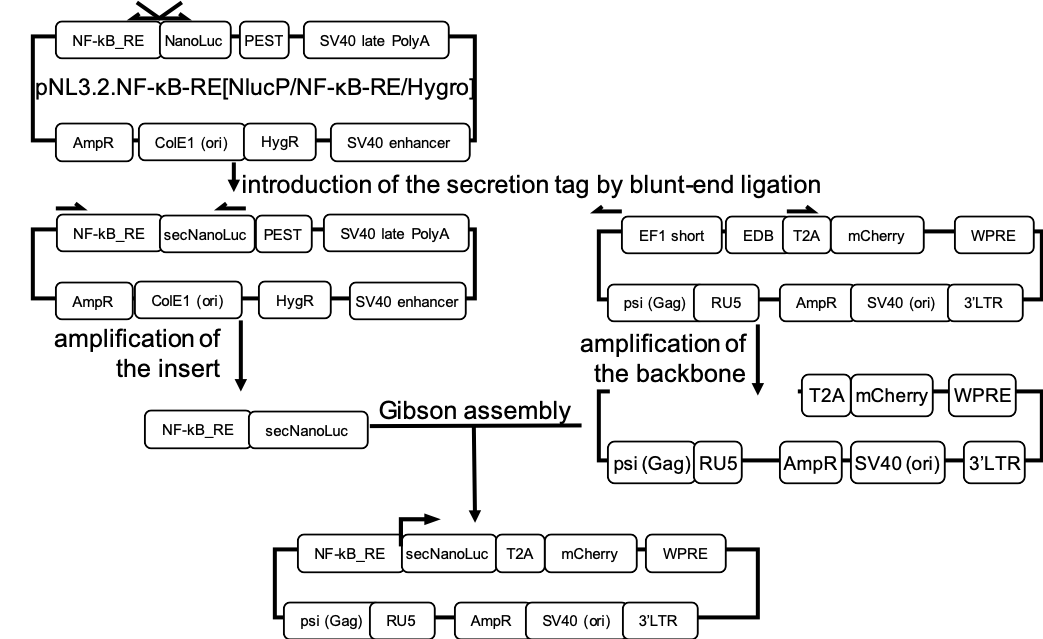 |
| --- | --- |
| **b** | NF-kB_RE-minimalPromoter-hIL6_secretionSignal-NanoLuc-T2A-mCherry |
|  | GGGAATTTCCGGGGACTTTCCGGGAATTTCCGGGGACTTTCCGGGAATTTCC-AGATCTGG CCTCGGCGGCCAAGCTTAGACACT-AGAGGGTATATAATGGAAGCTCGACTTCCAG-CTTG GCAATCCGGTACTGTTGGTAAAGCCACC-ATGAACTCCTTCTCCACAAGCGCCTTCGGTCC AGTTGCCTTCTCCCTGGGCCTGCTCCTGGTGTTGCCTGCTGCCTTCCCTGCCCCA-GTCTT CACACTCGAAGATTTCGTTGGGGACTGGCGACAGACAGCCGGCTACAACCTGGACCAAGTCCTTGAACAGGGAGGTGTGTCCAGTTTGTTTCAGAATCTCGGGGTGTCCGTAACTCCGATCCAAAGGATTGTCCTGAGCGGTGAAAATGGGCTGAAGATCGACATCCATGTCATCATCCCGTATGAAGGTCTGAGCGGCGACCAAATGGGCCAGATCGAAAAAATTTTTAAGGTGGTGTACCCTGTGGATGATCATCACTTTAAGGTGATCCTGCACTATGGCACACTGGTAATCGACGGGGTTACGCCGAACATGATCGACTATTTCGGACGGCCGTATGAAGGCATCGCCGTGTTCGACGGCAAAAAGATCACTGTAACAGGGACCCTGTGGAACGGCAACAAAATTATCGACGAGCGCCTGATCAACCCCGACGGCTCCCTGCTGTTCCGAGTAACCATCAACGGAGTGACCGGCTGGCGGCTGTGCGAACGCATTCTGGCG-GAGGGCAGAGGAAGTCTTCTAACATGCGGTGACGTGGAGGAGA ATCCCGGCCCT-ATGGTGAGCAAGGGCGAGGAGGATAACATGGCCATCATCAAGGAGTTCA TGCGCTTCAAGGTGCACATGGAGGGCTCCGTGAACGGCCACGAGTTCGAGATCGAGGGCGAGGGCGAGGGCCGCCCCTACGAGGGCACCCAGACCGCCAAGCTGAAGGTGACCAAGGGTGGCCCCCTGCCCTTCGCCTGGGACATCCTGTCCCCTCAGTTCATGTACGGCTCCAAGGCCTACGTGAAGCACCCCGCCGACATCCCCGACTACTTGAAGCTGTCCTTCCCCGAGGGCTTCAAGTGGGAGCGCGTGATGAACTTCGAGGACGGCGGCGTGGTGACCGTGACCCAGGACTCCTCCCTGCAGGACGGCGAGTTCATCTACAAGGTGAAGCTGCGCGGCACCAACTTCCCCTCCGACGGCCCCGTAATGCAGAAGAAGACCATGGGCTGGGAGGCCTCCTCCGAGCGGATGTACCCCGAGGACGGCGCCCTGAAGGGCGAGATCAAGCAGAGGCTGAAGCTGAAGGACGGCGGCCACTACGACGCTGAGGTCAAGACCACCTACAAGGCCAAGAAGCCCGTGCAGCTGCCCGGCGCCTACAACGTCAACATCAAGTTGGACATCACCTCCCACAACGAGGACTACACCATCGTGGAACAGTACGAACGCGCCGAGGGCCGCCACTCCACCGGCGGCATGGACGAGCTGTACAAGTAA |
|  | Supplementary Figure S1: cloning strategy and sequence of the reporter construct a) outline of the cloning strategy b) sequence of the reporter construct, primer binding sites are underlined |

RU5-Psi (Gag)-EF1(short)-**NF-kB_RE(response element)**-minimal promoter (minP)-IL6 secretion tag-Nanoluc-T2A-mCherry-WPRE-3’LTR-SV40 ori-Ampicillin resistance

GCGCGTTTCGGTGATGACGGTGAAAACCTCTGACACATGCAGCTCCCGGAGACGGTCACAGCTTGTCTGTAAGCGGATGCCGGGAGCAGACAAGCCCGTCAGGGCGCGTCAGCGGGTGTTGGCGGGTGTCGGGGCTGGCTTAACTATGCGGCATCAGAGCAGATTGTACTGAGAGTGCACCATATGCGGTGTGAAATACCGCACAGATGCGTAAGGAGAAAATACCGCATCAGGCGCCATTCGCCATTCAGGCTGCGCAACTGTTGGGAAGGGCGATCGGTGCGGGCCTCTTCGCTATTACGCCAGCTGGCGAAAGGGGGATGTGCTGCAAGGCGATTAAGTTGGGTAACGCCAGGGTTTTCCCAGTCACGACGTTGTAAAACGACGGCCAGTGCCAAGCTGACGCGTGTAGTCTTATGCAATACTCTTGTAGTCTTGCAACATGGTAACGATGAGTTAGCAACATGCCTTACAAGGAGAGAAAAAGCACCGTGCATGCCGATTGGTGGAAGTAAGGTGGTACGATCGTGCCTTATTAGGAAGGCAACAGACGGGTCTGACATGGATTGGACGAACCACTGAATTGCCGCATTGCAGAGATATTGTATTTAAGTGCCTAGCTCGATACAATAAACGGGTCTCTCTGGTTAGACCAGATCTGAGCCTGGGAGCTCTCTGGCTAACTAGGGAACCCACTGCTTAAGCCTCAATAAAGCTTGCCTTGAGTGCTTCAAGTAGTGTGTGCCCGTCTGTTGTGTGACTCTGGTAACTAGAGATCCCTCAGACCCTTTTAGTCAGTGTGGAAAATCTCTAGCAGTGGCGCCCGAACAGGGACCTGAAAGCGAAAGGGAAACCAGAGCTCTCTCGACGCAGGACTCGGCTTGCTGAAGCGCGCACGGCAAGAGGCGAGGGGCGGCGACTGGTGAGTACGCCAAAAATTTTGACTAGCGGAGGCTAGAAGGAGAGAGATGGGTGCGAGAGCGTCAGTATTAAGCGGGGGAGAATTAGATCGCGATGGGAAAAAATTCGGTTAAGGCCAGGGGGAAAGAAAAAATATAAATTAAAACATATAGTATGGGCAAGCAGGGAGCTAGAACGATTCGCAGTTAATCCTGGCCTGTTAGAAACATCAGAAGGCTGTAGACAAATACTGGGACAGCTACAACCATCCCTTCAGACAGGATCAGAAGAACTTAGATCATTATATAATACAGTAGCAACCCTCTATTGTGTGCATCAAAGGATAGAGATAAAAGACACCAAGGAAGCTTTAGACAAGATAGAGGAAGAGCAAAACAAAAGTAAGACCACCGCACAGCAAGCGGCCACTGATCTTCAGACCTGGAGGAGGAGATATGAGGGACAATTGGAGAAGTGAATTATATAAATATAAAGTAGTAAAAATTGAACCATTAGGAGTAGCACCCACCAAGGCAAAGAGAAGAGTGGTGCAGAGAGAAAAAAGAGCAGTGGGAATAGGAGCTTTGTTCCTTGGGTTCTTGGGAGCAGCAGGAAGCACTATGGGCGCAGCGTCAATGACGCTGACGGTACAGGCCAGACAATTATTGTCTGGTATAGTGCAGCAGCAGAACAATTTGCTGAGGGCTATTGAGGCGCAACAGCATCTGTTGCAACTCACAGTCTGGGGCATCAAGCAGCTCCAGGCAAGAATCCTGGCTGTGGAAAGATACCTAAAGGATCAACAGCTCCTGGGGATTTGGGGTTGCTCTGGAAAACTCATTTGCACCACTGCTGTGCCTTGGAATGCTAGTTGGAGTAATAAATCTCTGGAACAGATTTGGAATCACACGACCTGGATGGAGTGGGACAGAGAAATTAACAATTACACAAGCTTAATACACTCCTTAATTGAAGAATCGCAAAACCAGCAAGAAAAGAATGAACAAGAATTATTGGAATTAGATAAATGGGCAAGTTTGTGGAATTGGTTTAACATAACAAATTGGCTGTGGTATATAAAATTATTCATAATGATAGTAGGAGGCTTGGTAGGTTTAAGAATAGTTTTTGCTGTACTTTCTATAGTGAATAGAGTTAGGCAGGGATATTCACCATTATCGTTTCAGACCCACCTCCCAACCCCGAGGGGACCCGACAGGCCCGAAGGAATAGAAGAAGAAGGTGGAGAGAGAGACAGAGACAGATCCATTCGATTAGTGAACGGATCTCGACGGTATCGGTTAACTTTTAAAAGAAAAGGGGGGATTGGGGGGTACAGTGCAGGGGAAAGAATAGTAGACATAATAGCAACAGACATACAAACTAAAGAATTACAAAAACAAATTACAAAATTCAAAATTTTATCGATACTAGTGGATCTGCGATCGCAGGTGCCAGAACATTTCTCTGGCCTAACTGGCCGGTACCTGAGCTCGCTAGC**GGGAATTTCCGGGGACTTTCCGGGAATTTCCGGGGACTTTCCGGGAATTTCC**AGATCTGGCCTCGGCGGCCAAGCTTAGACACTAGAGGGTATATAATGGAAGCTCGACTTCCAGCTTGGCAATCCGGTACTGTTGGTAAAGCCACCATGAACTCCTTCTCCACAAGCGCCTTCGGTCCAGTTGCCTTCTCCCTGGGCCTGCTCCTGGTGTTGCCTGCTGCCTTCCCTGCCCCAGTCTTCACACTCGAAGATTTCGTTGGGGACTGGCGACAGACAGCCGGCTACAACCTGGACCAAGTCCTTGAACAGGGAGGTGTGTCCAGTTTGTTTCAGAATCTCGGGGTGTCCGTAACTCCGATCCAAAGGATTGTCCTGAGCGGTGAAAATGGGCTGAAGATCGACATCCATGTCATCATCCCGTATGAAGGTCTGAGCGGCGACCAAATGGGCCAGATCGAAAAAATTTTTAAGGTGGTGTACCCTGTGGATGATCATCACTTTAAGGTGATCCTGCACTATGGCACACTGGTAATCGACGGGGTTACGCCGAACATGATCGACTATTTCGGACGGCCGTATGAAGGCATCGCCGTGTTCGACGGCAAAAAGATCACTGTAACAGGGACCCTGTGGAACGGCAACAAAATTATCGACGAGCGCCTGATCAACCCCGACGGCTCCCTGCTGTTCCGAGTAACCATCAACGGAGTGACCGGCTGGCGGCTGTGCGAACGCATTCTGGCGGAGGGCAGAGGAAGTCTTCTAACATGCGGTGACGTGGAGGAGAATCCCGGCCCTATGGTGAGCAAGGGCGAGGAGGATAACATGGCCATCATCAAGGAGTTCATGCGCTTCAAGGTGCACATGGAGGGCTCCGTGAACGGCCACGAGTTCGAGATCGAGGGCGAGGGCGAGGGCCGCCCCTACGAGGGCACCCAGACCGCCAAGCTGAAGGTGACCAAGGGTGGCCCCCTGCCCTTCGCCTGGGACATCCTGTCCCCTCAGTTCATGTACGGCTCCAAGGCCTACGTGAAGCACCCCGCCGACATCCCCGACTACTTGAAGCTGTCCTTCCCCGAGGGCTTCAAGTGGGAGCGCGTGATGAACTTCGAGGACGGCGGCGTGGTGACCGTGACCCAGGACTCCTCCCTGCAGGACGGCGAGTTCATCTACAAGGTGAAGCTGCGCGGCACCAACTTCCCCTCCGACGGCCCCGTAATGCAGAAGAAGACCATGGGCTGGGAGGCCTCCTCCGAGCGGATGTACCCCGAGGACGGCGCCCTGAAGGGCGAGATCAAGCAGAGGCTGAAGCTGAAGGACGGCGGCCACTACGACGCTGAGGTCAAGACCACCTACAAGGCCAAGAAGCCCGTGCAGCTGCCCGGCGCCTACAACGTCAACATCAAGTTGGACATCACCTCCCACAACGAGGACTACACCATCGTGGAACAGTACGAACGCGCCGAGGGCCGCCACTCCACCGGCGGCATGGACGAGCTGTACAAGTAAGTCGACAATCAACCTCTGGATTACAAAATTTGTGAAAGATTGACTGGTATTCTTAACTATGTTGCTCCTTTTACGCTATGTGGATACGCTGCTTTAATGCCTTTGTATCATGCTATTGCTTCCCGTATGGCTTTCATTTTCTCCTCCTTGTATAAATCCTGGTTGCTGTCTCTTTATGAGGAGTTGTGGCCCGTTGTCAGGCAACGTGGCGTGGTGTGCACTGTGTTTGCTGACGCAACCCCCACTGGTTGGGGCATTGCCACCACCTGTCAGCTCCTTTCCGGGACTTTCGCTTTCCCCCTCCCTATTGCCACGGCGGAACTCATCGCCGCCTGCCTTGCCCGCTGCTGGACAGGGGCTCGGCTGTTGGGCACTGACAATTCCGTGGTGTTGTCGGGGAAATCATCGTCCTTTCCTTGGCTGCTCGCCTGTGTTGCCACCTGGATTCTGCGCGGGACGTCCTTCTGCTACGTCCCTTCGGCCCTCAATCCAGCGGACCTTCCTTCCCGCGGCCTGCTGCCGGCTCTGCGGCCTCTTCCGCGTCTTCGCCTTCGCCCTCAGACGAGTCGGATCTCCCTTTGGGCCGCCTCCCCGCCTGGTACCTTTAAGACCAATGACTTACAAGGCAGCTGTAGATCTTAGCCACTTTTTAAAAGAAAAGGGGGGACTGGAAGGGCTAATTCACTCCCAACGAAAATAAGATCTGCTTTTTGCTTGTACTGGGTCTCTCTGGTTAGACCAGATCTGAGCCTGGGAGCTCTCTGGCTAACTAGGGAACCCACTGCTTAAGCCTCAATAAAGCTTGCCTTGAGTGCTTCAAGTAGTGTGTGCCCGTCTGTTGTGTGACTCTGGTAACTAGAGATCCCTCAGACCCTTTTAGTCAGTGTGGAAAATCTCTAGCAGTAGTAGTTCATGTCATCTTATTATTCAGTATTTATAACTTGCAAAGAAATGAATATCAGAGAGTGAGAGGAACTTGTTTATTGCAGCTTATAATGGTTACAAATAAAGCAATAGCATCACAAATTTCACAAATAAAGCATTTTTTTCACTGCATTCTAGTTGTGGTTTGTCCAAACTCATCAATGTATCTTATCATGTCTGGCTCTAGCTATCCCGCCCCTAACTCCGCCCAGTTCCGCCCATTCTCCGCCCCATGGCTGACTAATTTTTTTTATTTATGCAGAGGCCGAGGCCGCCTCGGCCTCTGAGCTATTCCAGAAGTAGTGAGGAGGCTTTTTTGGAGGCCTAGACTTTTGCAGAGACGGCCCAAATTCGTAATCATGGTCATAGCTGTTTCCTGTGTGAAATTGTTATCCGCTCACAATTCCACACAACATACGAGCCGGAAGCATAAAGTGTAAAGCCTGGGGTGCCTAATGAGTGAGCTAACTCACATTAATTGCGTTGCGCTCACTGCCCGCTTTCCAGTCGGGAAACCTGTCGTGCCAGCTGCATTAATGAATCGGCCAACGCGCGGGGAGAGGCGGTTTGCGTATTGGGCGCTCTTCCGCTTCCTCGCTCACTGACTCGCTGCGCTCGGTCGTTCGGCTGCGGCGAGCGGTATCAGCTCACTCAAAGGCGGTAATACGGTTATCCACAGAATCAGGGGATAACGCAGGAAAGAACATGTGAGCAAAAGGCCAGCAAAAGGCCAGGAACCGTAAAAAGGCCGCGTTGCTGGCGTTTTTCCATAGGCTCCGCCCCCCTGACGAGCATCACAAAAATCGACGCTCAAGTCAGAGGTGGCGAAACCCGACAGGACTATAAAGATACCAGGCGTTTCCCCCTGGAAGCTCCCTCGTGCGCTCTCCTGTTCCGACCCTGCCGCTTACCGGATACCTGTCCGCCTTTCTCCCTTCGGGAAGCGTGGCGCTTTCTCATAGCTCACGCTGTAGGTATCTCAGTTCGGTGTAGGTCGTTCGCTCCAAGCTGGGCTGTGTGCACGAACCCCCCGTTCAGCCCGACCGCTGCGCCTTATCCGGTAACTATCGTCTTGAGTCCAACCCGGTAAGACACGACTTATCGCCACTGGCAGCAGCCACTGGTAACAGGATTAGCAGAGCGAGGTATGTAGGCGGTGCTACAGAGTTCTTGAAGTGGTGGCCTAACTACGGCTACACTAGAAGGACAGTATTTGGTATCTGCGCTCTGCTGAAGCCAGTTACCTTCGGAAAAAGAGTTGGTAGCTCTTGATCCGGCAAACAAACCACCGCTGGTAGCGGTGGTTTTTTTGTTTGCAAGCAGCAGATTACGCGCAGAAAAAAAGGATCTCAAGAAGATCCTTTGATCTTTTCTACGGGGTCTGACGCTCAGTGGAACGAAAACTCACGTTAAGGGATTTTGGTCATGAGATTATCAAAAAGGATCTTCACCTAGATCCTTTTAAATTAAAAATGAAGTTTTAAATCAATCTAAAGTATATATGAGTAAACTTGGTCTGACAGTTACCAATGCTTAATCAGTGAGGCACCTATCTCAGCGATCTGTCTATTTCGTTCATCCATAGTTGCCTGACTCCCCGTCGTGTAGATAACTACGATACGGGAGGGCTTACCATCTGGCCCCAGTGCTGCAATGATACCGCGAGACCCACGCTCACCGGCTCCAGATTTATCAGCAATAAACCAGCCAGCCGGAAGGGCCGAGCGCAGAAGTGGTCCTGCAACTTTATCCGCCTCCATCCAGTCTATTAATTGTTGCCGGGAAGCTAGAGTAAGTAGTTCGCCAGTTAATAGTTTGCGCAACGTTGTTGCCATTGCTACAGGCATCGTGGTGTCACGCTCGTCGTTTGGTATGGCTTCATTCAGCTCCGGTTCCCAACGATCAAGGCGAGTTACATGATCCCCCATGTTGTGCAAAAAAGCGGTTAGCTCCTTCGGTCCTCCGATCGTTGTCAGAAGTAAGTTGGCCGCAGTGTTATCACTCATGGTTATGGCAGCACTGCATAATTCTCTTACTGTCATGCCATCCGTAAGATGCTTTTCTGTGACTGGTGAGTACTCAACCAAGTCATTCTGAGAATAGTGTATGCGGCGACCGAGTTGCTCTTGCCCGGCGTCAATACGGGATAATACCGCGCCACATAGCAGAACTTTAAAAGTGCTCATCATTGGAAAACGTTCTTCGGGGCGAAAACTCTCAAGGATCTTACCGCTGTTGAGATCCAGTTCGATGTAACCCACTCGTGCACCCAACTGATCTTCAGCATCTTTTACTTTCACCAGCGTTTCTGGGTGAGCAAAAACAGGAAGGCAAAATGCCGCAAAAAAGGGAATAAGGGCGACACGGAAATGTTGAATACTCATACTCTTCCTTTTTCAATATTATTGAAGCATTTATCAGGGTTATTGTCTCATGAGCGGATACATATTTGAATGTATTTAGAAAAATAAACAAATAGGGGTTCCGCGCACATTTCCCCGAAAAGTGCCACCTGACGTCTAAGAAACCATTATTATCATGACATTAACCTATAAAAATAGGCGTATCACGAGGCCCTTTCGTCTC

Supplementary Figure S2: sequence of the lentiviral reporter plasmid pJM046; the important regions are highlighted in color

| **a** | **F8(dDb)-4-1BBL**: F8_VH-*linker*-F8_VL-*linker-*4-1BBL-*linker*-4-1BBL-*linker*-4-1BBL |
| --- | --- |
|  | EVQLLESGGGLVQPGGSLRLSCAASGFTFSLFTMSWVRQAPGKGLEWVSAISGSGGSTYYADSVKGRFTISRDNSKNTLYLQMNSLRAEDTAVYYCAKSTHLYLFDYWGQGTLVTVSS-*GGSGG*-EIVLTQS PGTLSLSPGERATLSCRASQSVSMPFLAWYQQKPGQAPRLLIYGASSRATGIPDRFSGSGSGTDFTLTISRLEPEDFAVYYCQQMRGRPPTFGQGTKVEIK-*SSSSGSSSSGSSSSG*-ATTQQGSPVFAKLLAKN QASLCNTTLNWHSQDGAGSSYLSQGLRYEEDKKELVVDSPGLYYVFLELKLSPTFTNTGHKVQGWVSLVLQAKPQVDDFDNLALTVELFPCSMENKLVDRSWSQLLLLKAGHRLSVGLRAYLHGAQDAYRDWELSYPNTTSFGLFLVKPDNPWE-*G*-ATTQQGSPVFAKLLAKNQASLCNTTLNWHSQDGAGSSY LSQGLRYEEDKKELVVDSPGLYYVFLELKLSPTFTNTGHKVQGWVSLVLQAKPQVDDFDNLALTVELFPCSMENKLVDRSWSQLLLLKAGHRLSVGLRAYLHGAQDAYRDWELSYPNTTSFGLFLVKPDNPWE-*G*-ATTQQGSPVFAKLLAKNQASLCNTTLNWHSQDGAGSSYLSQGLRYEEDKKELVVDSPGLY YVFLELKLSPTFTNTGHKVQGWVSLVLQAKPQVDDFDNLALTVELFPCSMENKLVDRSWSQLLLLKAGHRLSVGLRAYLHGAQDAYRDWELSYPNTTSFGLFLVKPDNPWE |
| **b** | **F8(scFv)-4-1BBL**: F8_VH-*linker*-F8_VL-*linker*-4-1BBL |
|  | EVQLLESGGGLVQPGGSLRLSCAASGFTFSLFTMSWVRQAPGKGLEWVSAISGSGGSTYYADSVKGRFTISRDNSKNTLYLQMNSLRAEDTAVYYCAKSTHLYLFDYWGQGTLVTVSS-*GGGGSGGGGSG GGG*-EIVLTQSPGTLSLSPGERATLSCRASQSVSMPFLAWYQQKPGQAPRLLIYGASSRATGIPDRF SGSGSGTDFTLTISRLEPEDFAVYYCQQMRGRPPTFGQGTKVEIK-*SSSSGSSSSGSSSS*-GATTQQG SPVFAKLLAKNQASLCNTTLNWHSQDGAGSSYLSQGLRYEEDKKELVVDSPGLYYVFLELKLSPTFTNTGHKVQGWVSLVLQAKPQVDDFDNLALTVELFPCSMENKLVDRSWSQLLLLKAGHRLSVGLRALHGAQDAYRDWELSYPNTTSFGLFLVKPDNPWE |
| **c** | **F8(dDb)-GITRL**: F8_VH-*linker*-F8_VL-*linker-*GITRL-*linker*-GITRL-*linker*-GITRL |
|  | EVQLLESGGGLVQPGGSLRLSCAASGFTFSLFTMSWVRQAPGKGLEWVSAISGSGGSTYYADSVKGRFTISRDNSKNTLYLQMNSLRAEDTAVYYCAKSTHLYLFDYWGQGTLVTVSS-*GGSGG*-EIVLTQS PGTLSLSPGERATLSCRASQSVSMPFLAWYQQKPGQAPRLLIYGASSRATGIPDRFSGSGSGTDFTLTISRLEPEDFAVYYCQQMRGRPPTFGQGTKVEIK-*SSSSGSSSSGSSSSG*-PTAIESCMVKFELSSSKW HMTSPKPHCVNTTSDGKLKILQSGTYLIYGQVIPVDKKYIKDNAPFVVQIYKKNDVLQTLMNDFQILPIGGVYELHAGDNIYLKFNSKDHIQKNNTYWGIILMPDLP-*GGGSGGG*-PTAIESCMVKFELSSSKW HMTSPKPHCVNTTSDGKLKILQSGTYLIYGQVIPVDKKYIKDNAPFVVQIYKKNDVLQTLMNDFQILPIGGVYELHAGDNIYLKFNSKDHIQKNNTYWGIILMPDLP-*GGGSGGG*-PTAIESCMVKFELSSSKW HMTSPKPHCVNTTSDGKLKILQSGTYLIYGQVIPVDKKYIKDNAPFVVQIYKKNDVLQTLMNDFQILPIGGVYELHAGDNIYLKFNSKDHIQKNNTYWGIILMPDLP |
| **d** | **F8(scFv)-GITRL**: F8_VH-*linker*-F8_VL-*linker*-GITRL |
|  | EVQLLESGGGLVQPGGSLRLSCAASGFTFSLFTMSWVRQAPGKGLEWVSAISGSGGSTYYADSVKGRFTISRDNSKNTLYLQMNSLRAEDTAVYYCAKSTHLYLFDYWGQGTLVTVSS-*GGGGSGGGGSG GGG*-EIVLTQSPGTLSLSPGERATLSCRASQSVSMPFLAWYQQKPGQAPRLLIYGASSRATGIPDRF SGSGSGTDFTLTISRLEPEDFAVYYCQQMRGRPPTFGQGTKVEIK-*SSSSGSSSSGSSSS*-PTAIESCM VKFELSSSKWHMTSPKPHCVNTTSDGKLKILQSGTYLIYGQVIPVDKKYIKDNAPFVVQIYKKNDVLQTLMNDFQILPIGGVYELHAGDNIYLKFNSKDHIQKNNTYWGIILMPDLP |
| **e** | **F8(scDb)-CD154**: F8_VH-*linker*-F8_VL- *linker*- F8_VH-*linker*-F8_VL-*linker*-CD154-*linker*-CD154-*linker*-CD154 |
|  | EVQLLESGGGLVQPGGSLRLSCAASGFTFSLFTMSWVRQAPGKGLEWVSAISGSGGSTYYADSVKGRFTISRDNSKNTLYLQMNSLRAEDTAVYYCAKSTHLYLFDYWGQGTLVTVSS-*GGSGG*-EIVLTQS PGTLSLSPGERATLSCRASQSVSMPFLAWYQQKPGQAPRLLIYGASSRATGIPDRFSGSGSGTDFTLTISRLEPEDFAVYYCQQMRGRPPTFGQGTKVEIK-*GGGGSGGGGSGGGGS*-EVQLLESGGGLVQP GGSLRLSCAASGFTFSLFTMSWVRQAPGKGLEWVSAISGSGGSTYYADSVKGRFTISRDNSKNTLYLQMNSLRAEDTAVYYCAKSTHLYLFDYWGQGTLVTVSS-*GGSGG*-EIVLTQSPGTLSLSPGERATLS CRASQSVSMPFLAWYQQKPGQAPRLLIYGASSRATGIPDRFSGSGSGTDFTLTISRLEPEDFAVYYCQQMRGRPPTFGQGTKVEIK-*SSSSGSSSSGSSSS*-QRGDEDPQIAAHVVSEANSNAASVLQWAKK GYYTMKSNLVMLENGKQLTVKREGLYYVYTQVTFCSNREPSSQRPFIVGLWLKPSSGSERILLKANTHSSSQLCEQQSVHLGGVFELQAGASVFVNVTEASQVIHRVGFSSFGLLKL-*GGGS*-QRGDEDPQIA AHVVSEANSNAASVLQWAKKGYYTMKSNLVMLENGKQLTVKREGLYYVYTQVTFCSNREPSSQRPFIVGLWLKPSSGSERILLKANTHSSSQLCEQQSVHLGGVFELQAGASVFVNVTEASQVIHRVGFSSFGLLKL-*GGGS*-QRGDEDPQIAAHVVSEANSNAASVLQWAKKGYYTMKSNLVMLENGKQLTVKR EGLYYVYTQVTFCSNREPSSQRPFIVGLWLKPSSGSERILLKANTHSSSQLCEQQSVHLGGVFELQAGASVFVNVTEASQVIHRVGFSSFGLLKL |
| Supplementary Figure S3: Sequences of the immunocytokines that were developed in this study, the linker sequences are depicted in italics | |

**SUPPLEMENTARY MATERIALS AND METHODS**

SDS PAGE

The proteins were diluted to 0.1 mg/mL and 4 μL Lämmli buffer (0.25 M Tris-HCl pH 6.8, 40% glycerol, 8% sodium dodecyl sulfate, 0.02% bromophenol blue) was added to 12 μL of the protein dilution. Reducing Lämmli buffer was prepared fresh by adding 20% β-mercaptoethanol to the non-reducing Lämmli buffer. The samples were incubated at 95°C for 5 min and then directly loaded onto a TruPAGE Precasd 4-12% gel (Sigma Aldrich, #PCG2003 or #PCG2007). The gel was run in TruPAGE TEA-Tricine buffer (Sigma Aldrich, #PCG3001) diluted according to the manufacturer’s recommendations at 180 V, 110 mAmp for 1 h. Afterwards, the gel was rinsed with deionized water and stained for 1 h in Coomassie blue (2.4 g/L Brilliant blue, 24% methanol, 8% acetic acid). The gel was then destained in destaining solution (30% methanol, 10% acetic acid) for several hours at room temperature. Finally, the gel was left in deionized water at room temperature overnight to complete the destaining process.

Size exclusion chromatography

For size exclusion chromatography, a Superdex 200 10/300 increase column (GE healthcare) was used together with an Äkta Pure FPLC machine (GE healthcare). The size exclusion chromatography was run in PBS at a speed of 0.75 mL/min.

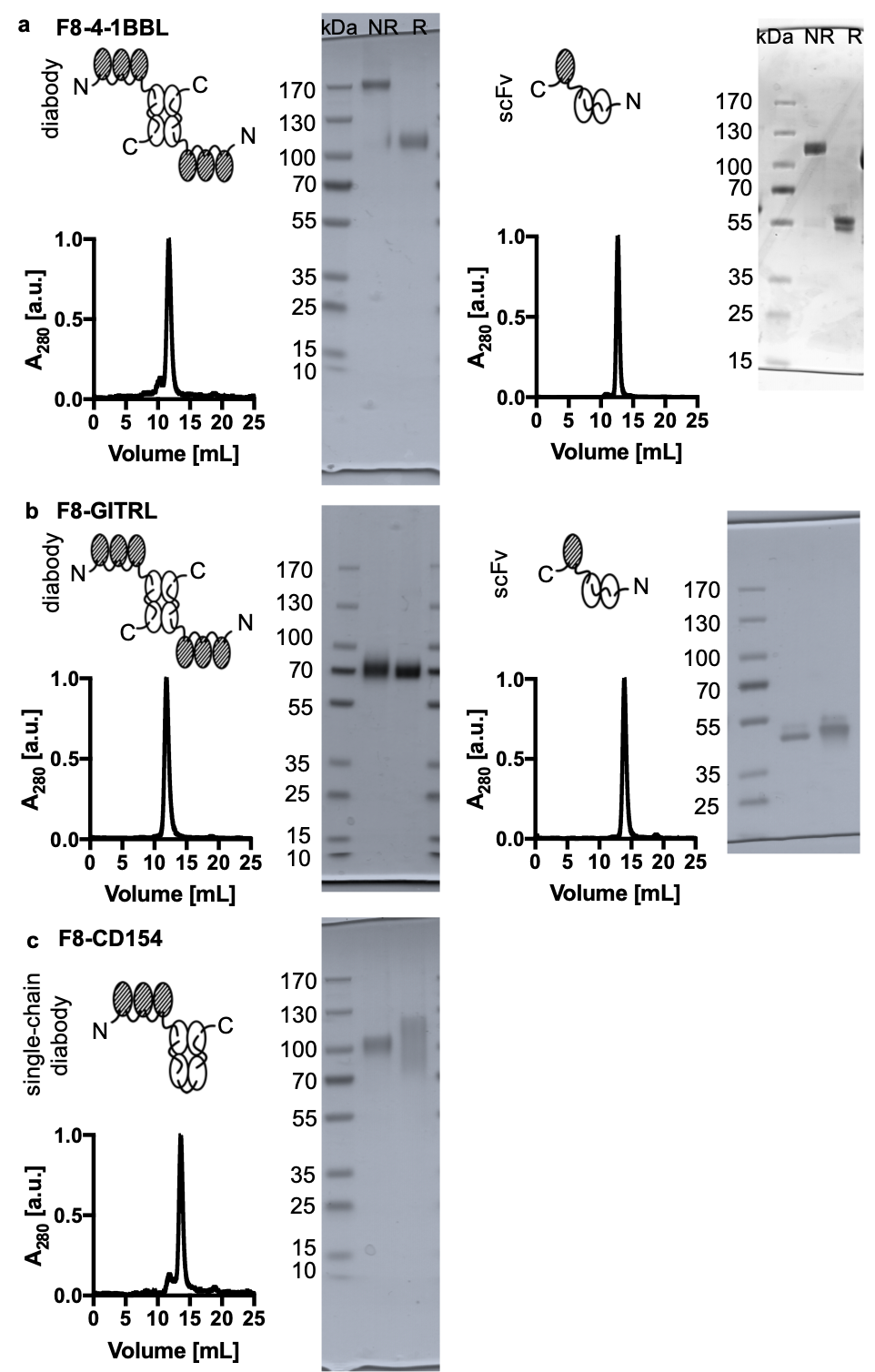

Supplementary Figure S4: The proteins were characterized by SDS PAGE (NR: non-reducing, R: reducing sample buffer) and size exclusion chromatography (Superdex 200 Increase, 10/300 GL, GE Healthcare) after purification.

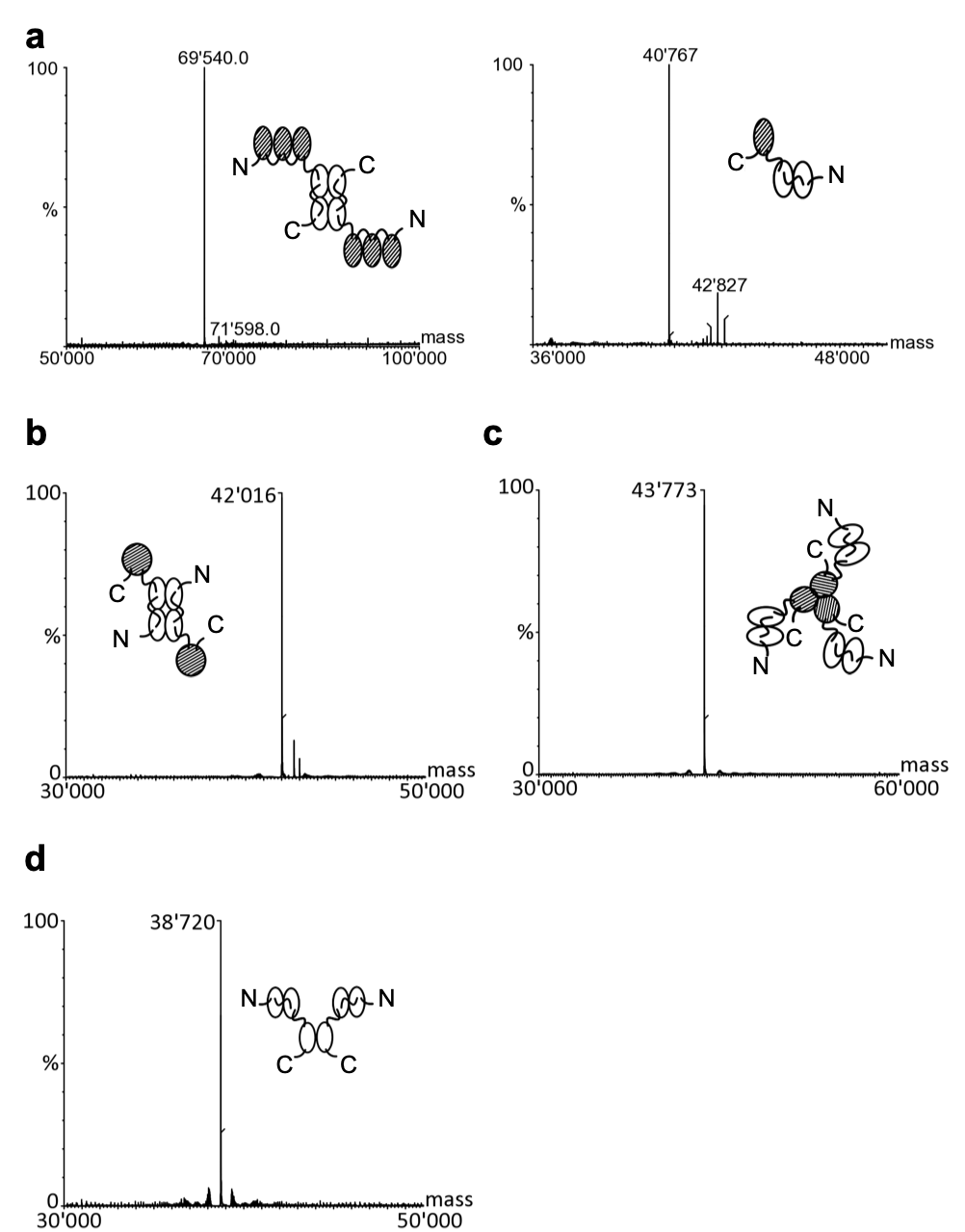

Supplementary Figure S5: Mass spectrometry data for a) the F8-GITRL constructs, b) L19-IL2, c) F8-TNF and d) F8 in the small immune protein (sip) format

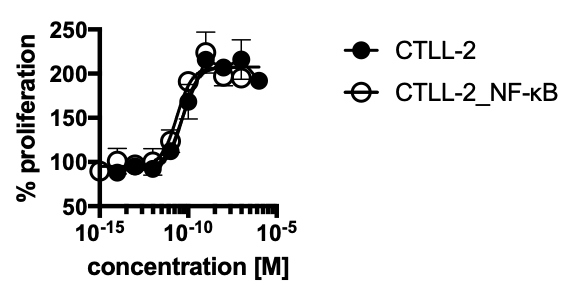

Supplementary Figure S6: Comparison of the conventional CTLL-2 proliferation both with the transduced and the non-transduced cell line using L19-IL2

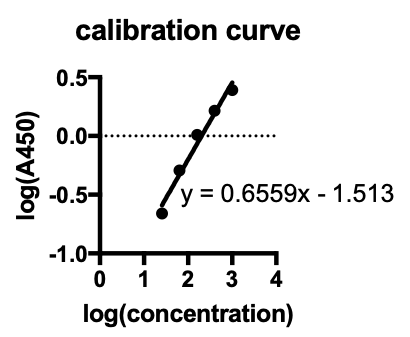

Supplementary Figure S7: Standard curve used to determine the IFN-γ concentration from the A450 measurements. A linear curve fit was applied using the GraphPad Prism 7.0 a software and this curve fit was then used to calculate the IFN-γ concentration from the absorbance measurements.

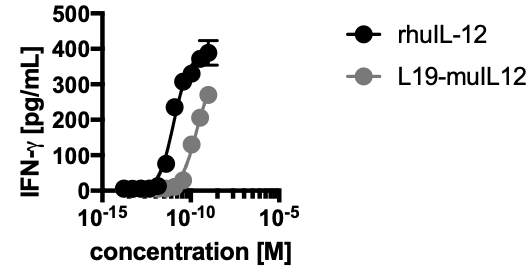

Supplementary Figure S8: The standard activity of IL-12 measuring the IFN-g release by NK-92 cells was performed both with the L19-IL12 immunocytokine featuring the murine IL-12 and with WHO calibrated recombinant human IL-12. Since IL-12 is not 100% cross-reactive between mouse and human, differences in activity were observed.

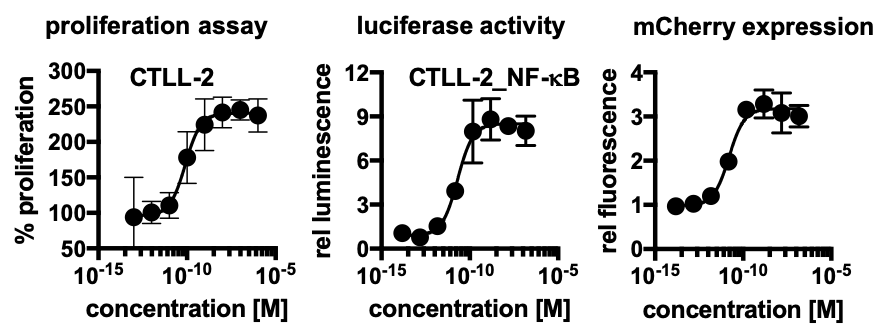

Supplementary Figure S9: The CTLL-2 proliferation assay as well as the NF-κB reporter assays were also performed with recombinant human IL-2 (Proleukin, Roche Diagnostics)

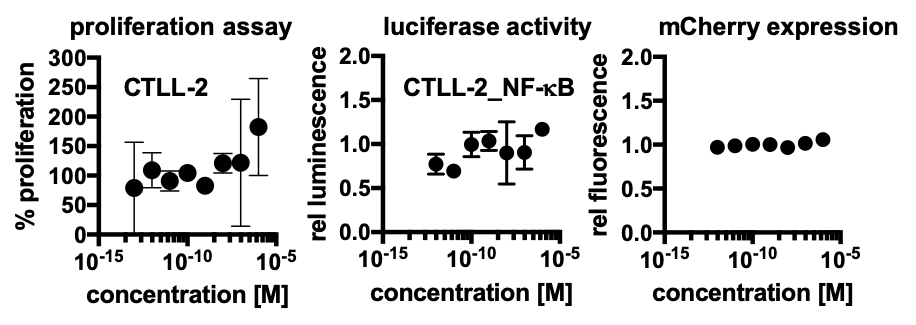

Supplementary Figure S10: The CTLL-2 proliferation assay as well as the NF-κB reporter assays were also performed with “naked” F8 in the small immune protein format as a negative control

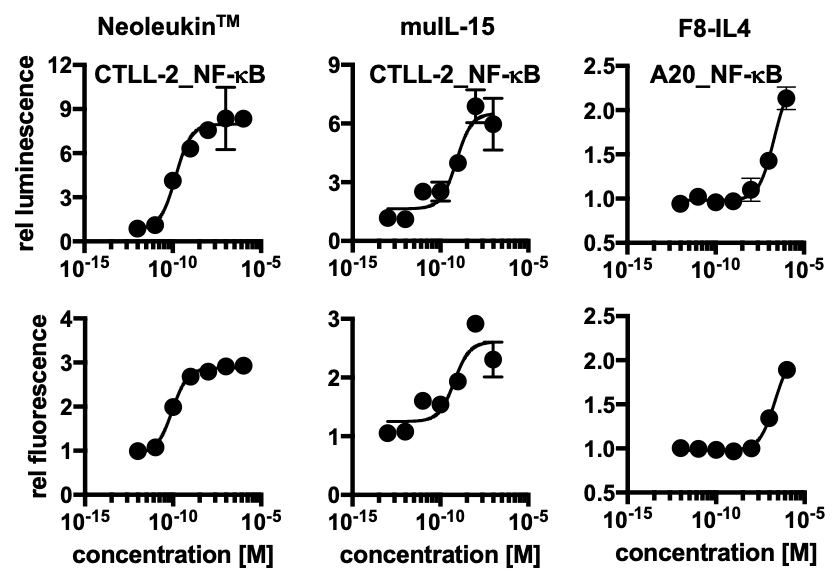

Supplementary Figure S11: The NF-kB reporter assay were also performed with recombinant Neoleukin^TM^, a fully synthetic cytokine that was first described by Silva *et al.^1^*, recombinant murine IL-15 (Peprotech, #210-15) and the F8-IL4 fusion protein^2^

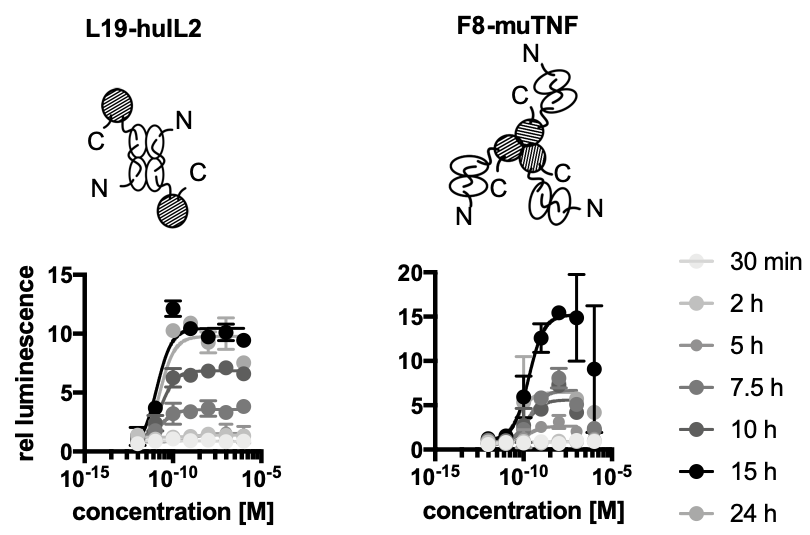

Supplementary Figure S12: The luciferase assay was performed at different time points after addition of the cytokines. In both cases, the signal was strongest after 15 h which corresponds to the time point when the samples were taken for the other measurements.

|  | **conventional assay** | | | **luciferase assay** | | | **mCherry expression** | | |
| --- | --- | --- | --- | --- | --- | --- | --- | --- | --- |
|  | **EC_50_** | **95% CI** | **R^2^** | **EC_50_** | **95% CI** | **R^2^** | **EC_50_** | **95% CI** | **R^2^** |
| L19-IL2 | 47 pM | 26 to 83 pM | 0.95 | 15 pM | 8.5 to 26 pM | 0.94 | 21 pM | 6.3 to 74 pM | 0.82 |
| L19-IL12 | 0.1 nM | 0.1 to 0.2 nM | 0.99 | 3.5 nM | 1.9 to 6.0 nM | 0.95 | 2.6 nM | 2.0 to 3.4 nM | 0.99 |
| F8-TNF | 0.7 pM | 0.6 to 0.9 pM | 0.98 | 450 pM | 280 to 730 pM | 0.96 | 540 pM | 450 to 650 pM | 0.99 |

Supplementary Table T1: EC_50_ obtained from a sigmoidal curve fit for conventional assay as well as for the readouts with the new cell lines (95% CI: 95% confidence interval)

|  |  | **+ EDA** | | | | | | **no EDA** | | | | | |
| --- | --- | --- | --- | --- | --- | --- | --- | --- | --- | --- | --- | --- | --- |
|  |  | **luciferase activity** | | | **mCherry expression** | | | **luciferase activity** | | | **mCherry expression** | | |
|  |  | **EC_50_** | **95% CI** | **R^2^** | **EC_50_** | **95% CI** | **R^2^** | **EC_50_** | **95% CI** | **R^2^** | **EC_50_** | **95% CI** | **R^2^** |
| F8-4-1BBL | diabody | 1.1 nM | 0.9 to 1.4 nM | 0.99 | 1.2 nM | 0.9 to 1.5 nM | 0.99 | 1.1 nM | 0.8 to 1.5 nM | 0.98 | 1.0 nM | 0.8 to 1.3 nM | 0.99 |
|  | scFv | 1.1 nM | 0.7 to 1.8 nM | 0.97 | 1.0 nM | 0.7 to 1.6 nM | 0.98 | n/a | n/a | n/a | n/a | n/a | n/a |
| F8-GITRL | diabody | 7.6 nM | 6.2 to 9.3 nM | 0.99 | 7.9 nM | 7.0 to 8.9 nM | 1.00 | 4.7 nM | 3.9 to 5.7 nM | 0.99 | 4.1 nM | 3.6 to 4.6 nM | 1.00 |
|  | scFv | 11 nM | 7.5 to 18 nM | 0.97 | 9.6 nM | 7.5 to 12 nM | 0.99 | 56 nM | 43 to 72 nM | 0.99 | 67 nM | 57 to 79 nM | 0.99 |
| F8-CD154 | diabody | 65 pM | 45 to 9.3 pM | 0.98 | 64 pM | 49 to 85 pM | 0.99 | 660 pM | 0.4 to 1.1 nM | 0.97 | 1.1 nM | 0.4 to 2.7 nM | 0.93 |

Supplementary Table T2: EC_50_ values obtained from a sigmoidal curve fit for the different F8-TNFSF fusion proteins (95% CI: 95% confidence interval)

1	Silva, D. A. *et al.* De novo design of potent and selective mimics of IL-2 and IL-15. *Nature* **565**, 186-191, doi:10.1038/s41586-018-0830-7 (2019).

2	Hemmerle, T. & Neri, D. The antibody-based targeted delivery of interleukin-4 and 12 to the tumor neovasculature eradicates tumors in three mouse models of cancer. *Int J Cancer* **134**, 467-477, doi:10.1002/ijc.28359 (2014).
